# Integrative genomic analysis unifying epigenetic inheritance in adaptation and canalization

**DOI:** 10.1101/849620

**Authors:** Abhay Sharma

## Abstract

Epigenetic inheritance, especially its biomedical and evolutionary significance, is an immensely interesting but highly controversial subject. Notably, a recent analysis of existing multi-omics has supported the mechanistic plausibility of epigenetic inheritance and its implications in disease and evolution. The evolutionary support stemmed from the specific finding that genes associated with cold induced inheritance and with latitudinal adaptation in mice are exceptionally common. Here, a similar gene set overlap analysis is presented that integrates cold induced inheritance with evolutionary adaptation and genetic canalization in cold environment in *Drosophila*. Genes showing differential expression in inheritance specifically overrepresent gene sets associated with differential and allele specific expression, though not with genome-wide genetic differentiation, in adaptation. On the other hand, the differentiated outliers uniquely overrepresent genes dysregulated by radicicol, a decanalization inducer. Both gene sets in turn exclusively show enrichment of genes that accumulate, in intended experimental lines, *de novo* mutations, a potential source of canalization. Successively, the three gene sets distinctively overrepresent genes exhibiting, between mutation accumulation lines, invariable expression, a potential signal for canalization. Sequentially, the four gene sets solely display enrichment of genes grouped in gene ontology under transcription factor activity, a signature of regulatory canalization. Cumulatively, the analysis suggests that epigenetic inheritance possibly contributes to evolutionary adaptation in the form of *cis* regulatory variations, with *trans* variations arising in the course of genetic canalization.

## Introduction

Recent evidence supporting environmentally induced germline epigenetic inheritance in diverse species has sparked immense interest and controversy especially over its potential evolutionary and disease significance (Li *et al.*, 2012; Horsthemke, 2018; Radford, 2018; Skvortsova *et al.*, 2018; Anastasiadi and Piferrer, 2019; Cavalli and Heard, 2019; Lesch *et al*., 2019; Perez and Lehner, 2019). Integrative genomics is considered as a valuable approach in addressing these advanced prospects of epigenetic inheritance (Molaro *et al.*, 2011; Sharma, 2015; Cavalli and Heard, 2019). Indeed, a multi-omics data analysis has recently revealed an association between cold induced inheritance and latitudinal adaptation in mice, and between epigenetic inheritance and disease in humans (Sharma, 2019). In mice, genes that are differentially methylated in sperm inheriting cold induced traits have been found to specifically overrepresent genes related to latitudinal adaptation associated single nucleotide polymorphisms, and differential and allele specific expression (Sharma, 2019). Similarly, in humans, differentially methylated genes in sperm of enriched-risk cohort of fathers of autistic children specifically overrepresent genes harbouring *de novo* mutations in autism (Sharma, 2019). Also, differentially methylated genes in sperm of recreational cannabis users specifically overrepresent genes harbouring *de novo* mutations in autism and schizophrenia, with paternal cannabis being a known suspect in psychotic disorders (Sharma, 2019). Notably, differentially methylated sperm genes and disease associated genes in general are enriched for genes known to exhibit high mutability and loss of function intolerance, suggesting a mechanistic basis for disease consequences of epigenetic inheritance (Sharma, 2019). Lastly, supporting developmental mechanisms of epigenetic inheritance in both mice and humans, the sperm genes in these species were found to overrepresent genes that are known to escape demethylation reprogramming and to express across stages in development (Sharma, 2019). Here, to further examine the evolutionary potential of epigenetic inheritance, a similar integrative genomic analysis of cold induced inheritance and latitudinal adaptation in the fruit fly *Drosophila melanogaster* is presented.

## Methods

As in mice and humans (Sharma, 2019), the present analysis in *D. melanogaster* is based on hypergeometric test of overlap between diverse gene sets, with the latter consisting of existing data pertaining to whole fly genome, differential gene expression, allele specific expression, single nucleotide polymorphism association, *de novo* mutation, rare and common genetic variation, and gene ontology. The original sources of gene sets used, along with remarks and relation to various figure panels shown, are detailed in **Supplementary Table 1**. Literature-wide search was undertaken to identify relevant studies. To remove any bias, only originally provided, not self defined, datasets were used. Standard gene symbols were used (Lyne et al., 2007). Gene set overlaps found are expressed as log_2_ of fold change, a positive value indicating overrepresentation or enrichment and a negative value indicating underrepresentation or depletion. Hypergeometric *p* value was ignored because the hypothesis testing primarily depended on comparison of fold change magnitude. Overlapping genes related to results presented in various figure panels are listed in **Supplementary Table 2**.

## Results and discussion

Before attempting to examine in details the impact of epigenetic inheritance on adaptive evolution, the specificity of the former with respect to environmental factor that induces inheritance and that underlies adaptation was tested first as background check. A total of three inheritance models were available for analysis, with transcriptomic effects of paternal F0 exposure reported in both F1 and F2 in two models (Sharma and Singh, 2009; Mohammad *et al.*, 2009; Teltumbade *et al.*, 2019) and in F1 alone in one study (Zare *et al.*, 2018). The environmental factors associated with these three models included high-sugar diet (HSD), the neuroactive compound pentylenetetrazole (PTZ), and cold exposure (CE). Though F2 phenotype is minimally required to demonstrate germline mediated inheritance, the study that was restricted to F1 only, the CE model, was also included in the analysis since biochemical level F2 effect has been demonstrated (Karunakar *et al.*, 2019) in CE induced inheritance. For uniformity, differentially expressed genes in parent (F0) and offspring (F1) were used in the analysis. The null hypothesis initially tested was that the magnitude of gene overlap between F0 and F1 in a given model is the same as that between F0 in that model and F1 in all the three models combined. Alternatively, the overlap between the former is higher than the latter, supporting environmental specificity of inherited expression phenotype. Notably, in general, a higher overlap was observed between F0 and F1 of a given model (Fig. 1a). This demonstrated that offspring inherit environmental factor specific gene regulatory changes. Next, environmental specificity between inheritance and adaptation was examined. The null hypothesis was similarly tested by replacing F0 samples with cold adaptive genes. The adaptation gene sets (Zhao *et al.*, 2015; Juneja *et al.*, 2016; Pool *et al.*, 2017; Mateo *et al.*, 2018; Kolaczkowski *et al.*, 2011) comprised of differentially expressed genes (DEG-LA), allele-specific expression (ASE-LA), and genome-wide single nucleotide polymorphism association (SNP-LA). Interestingly, DEG-LA and ASE-LA, not SNP-LA, were found to enrich only CE-F1, not HSD-F1 and PTZ-F1 (Fig. 1b). As ASE-LA reflects *cis* regulatory genetic variations, a lack of CE-F1 enrichment in SNP-LA suggested that the latter may signify *trans* acting regulatory adaptation, with the latter known as a major signature of gene expression canalization in cold adaptation in fly (von Heckel et al., 2016).

**Fig. 1.**
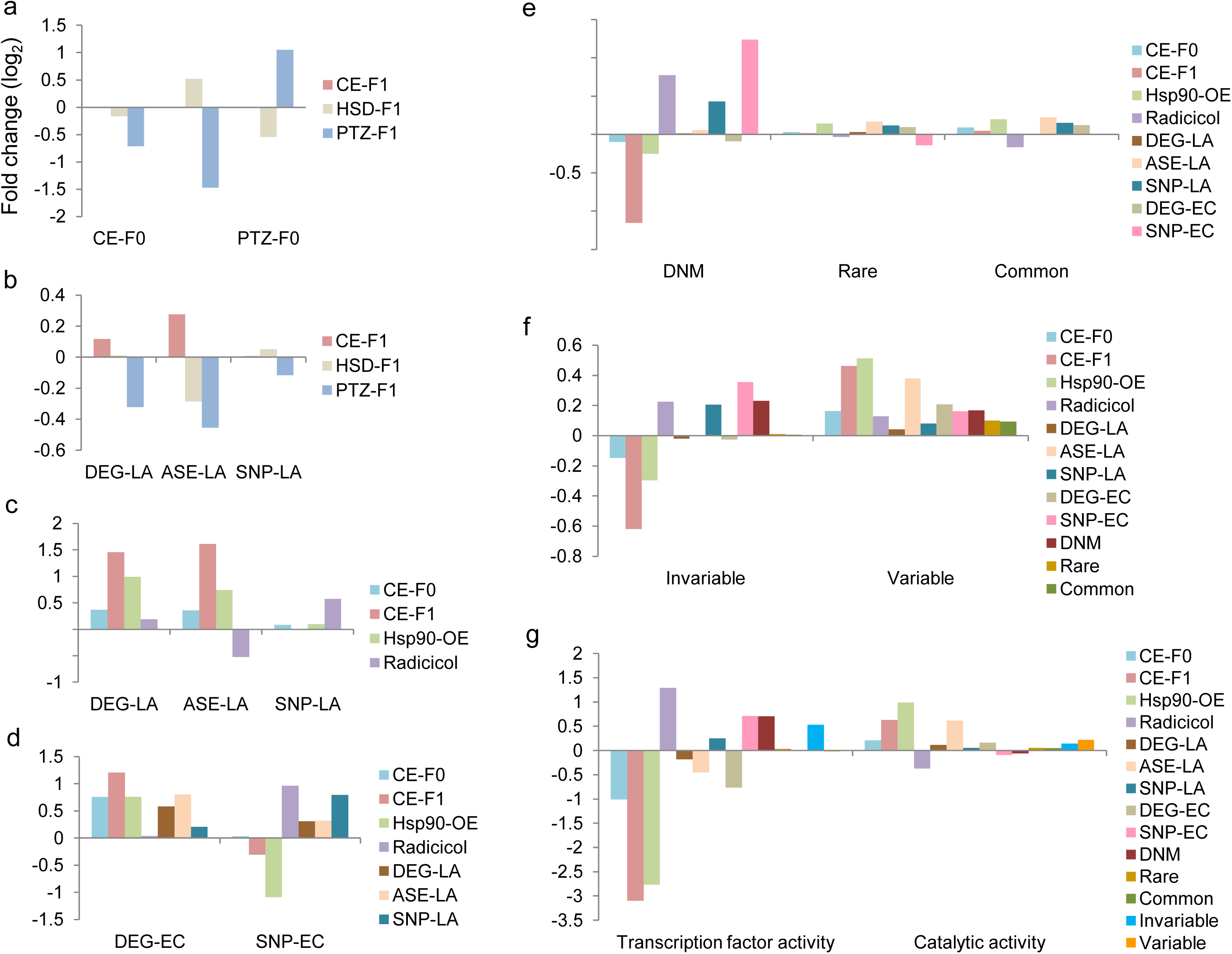
Graphical display of multi-omics gene set overlaps. Fold enrichment or depletion of (**a**) offspring genes in parental gene sets, (**b**) offspring genes in latitudinal adaptation gene sets, (**c**) offspring or hsp90 related genes in latitudinal adaptation gene sets, (**d**) the preceding panel’s gene sets in microclimatic adaptation genes, (**e**) the preceding panel’s gene sets in *de novo* mutation, and rare and common variation containing genes, (f) the preceding panel’s gene sets in invariable and variable expression associated genes in *de novo* mutation accumulation lines, and (g) the preceding panel’s gene sets in gene ontology genes grouped under the specified molecular function categories. Offspring genes combined (a, b) or whole genome (c-g) was used as background population in overlap analysis. Abbreviations and other details as explained in text. Exact source of gene lists used along with remarks and reference to figure panels are indicated in **Supplementary Table 1**. Overlapping genes are listed in **Supplementary Table 2**.

Next, I focused on cold inheritance model to investigate evolutionary implications of epigenetic inheritance in details. The hypergeometric tests performed henceforth in the analysis used genes in the whole genome as total population. Also, to remain unbiased, samples analyzed in each test were carried forward to the next. First, gene overlap test was performed with fly genes that are differentially expressed following treatment with radicicol, an inhibitor of Hsp90 that regulates canalization via diverse modes (Sawarkar *et al.*, 2012; Sawarkar and Paro, 2013). CE-F0 and genes differentially expressed in flies overexpressing Hsp90 (Hsp90-OE) were included in the test as controls (Ray *et al.*, 2019). As expected from background check (Fig. 1b), CE-F1 was overrepresented in DEG-LA, and ASE-LA, not SNP-LA. Remarkably, a clear overrepresentation of radicicol, not CE-F0, CE-F1, or Hsp90-OE, was observed in SNP-LA (Fig. 1c). Thereafter, all the above samples were challenged with gene sets associated with adaptation in Evolution Canyon, a microclimatic gradient of temperature, besides others (Yablonovitch *et al.*, 2017). The Evolution Canyon sets included genes with differentially expression (DEG-EC) and genome-wide single nucleotide polymorphism association (SNP-EC). Notably, radicicol and SNP-LA were found to be most highly enriched in SNP-EC (Fig. 1d). In contrast, CE-F1 was most highly enriched in DEG-EC. These results overall suggested that genome-wide association signals in evolutionary adaptation mostly relate to *trans* variations involved in regulatory canalization. This is consistent with one or more orders of magnitude larger mutational target space for *trans* than *cis* effects, and hence greater scope for *trans* variations in canalization of gene expression (Ehrenreich and Pfennig, 2016).

As *de novo* mutations are considered to act as source of variations in canalization (Fanti *et al.*, 2017), I next tested enrichment of an available compendium of *de novo* mutations associated genes (DNM) assembled from various mutation accumulation lines, with rare and common genetic variations as controls (Assaf *et al.*, 2017). Interestingly, radicicol, SNP-LA, and SNA-EC, not others, showed explicit overrepresentation of DNM (Fig. 1e). No such trend was observed with respect to rare or common variations. Further, given that gene expression between DNM lines can be invariable or variable possibly signifying canalization or uncanalization, in that order (Rifkin *et al.*, 2005), the samples were challenged with the invariable and variable gene sets. Remarkably, radicicol, SNP-LA, SNA-EC, and DNM were found to exclusively overrepresent invariable genes (Fig. 1f). In contrast, CE-F1, besides others, most prominently overrepresented variable genes. Finally, considering reported excess of transcription regulators and enzymes in genes with lower or higher expression variation, in that order, in DNM lines (Rifkin *et al.*, 2005), with transcription factors as most direct *trans* acting factor in canalization (Ehrenreich and Pfennig, 2016), the overlap test was next performed with genes grouped in gene ontology molecular function categories transcription factor activity and catalytic activity (Ashburner *et al.*, 2000; Carbon *et al.*, 2009). Indeed, radicicol, SNP-LA, SNA-EC, DNM, and Invariable genes were exclusively found enriched for transcription factor activity (Fig. 1g). Also, as expected, CE-F1, besides others, was enriched for catalytic activity. This completely unbiased gene ontology evidence independently suggested excess of *trans* regulatory variations in differentiated genomic outliers.

Over and above the recent multi-omics evidence broadly implicating epigenetic inheritance in evolutionary adaptation in mice, the present analysis specifically suggests that inheritance of acquired traits may possibly drive adaptation by contributing *cis* regulatory variations. In-depth studies are now required to confirm these albeit oversimplified initial observations. With expanding evidence of epigenetic inheritance, the call has increased for accommodating the latter in the modern evolutionary theory. The present results providing unbiased multi-omics evidence strongly supports this call.

## Supplementary materials

Supplementary Table 1. Details of gene sets used in the analysis.

Supplementary Table 2. List of overlapping genes.

## References

1. Ashburner M, Ball CA, Blake JA, Botstein D, Butler H, Cherry JM, Davis AP, Dolinski K, Dwight SS, Eppig JT, Harris MA, Hill DP, Issel-Tarver L, Kasarskis A, Lewis S, Matese JC, Richardson JE, Ringwald M, Rubin GM, Sherlock G. Gene ontology: tool for the unification of biology. The Gene Ontology Consortium. Nat. Genet. 25, 25–29 (2000).

2. Anastasiadi, D. & Piferrer, F. Epimutations in developmental genes underlie the onset of domestication in farmed European sea bass. Mol. Biol. Evol. doi: 10.1093/molbev/msz153 (2019).

3. Assaf ZJ, Tilk S, Park J, Siegal ML, Petrov DA. Deep sequencing of natural and experimental populations of Drosophila melanogaster reveals biases in the spectrum of new mutations. Genome Res. 27, 1988–2000 (2017).

4. Carbon S, Ireland A, Mungall CJ, Shu S, Marshall B, Lewis S; AmiGO Hub; Web Presence Working Group. AmiGO: online access to ontology and annotation data. Bioinformatics 25, 288–289 (2009).

5. Cavalli, G. & Heard, E. Advances in epigenetics link genetics to the environment and disease. Nature 571, 489–499 (2019).

6. Ehrenreich IM, Pfennig DW. Genetic assimilation: a review of its potential proximate causes and evolutionary consequences. Ann. Bot. 117, 769–779 (2016).

7. Fanti L, Piacentini L, Cappucci U, Casale AM, Pimpinelli S. Canalization by Selection of de Novo Induced Mutations. Genetics 206, 1995–2006 (2017).

8. Horsthemke, B. A critical view on transgenerational epigenetic inheritance in humans. Nat. Commun. 9, 2973 (2018).

9. Juneja P, Quinn A, Jiggins FM. Latitudinal clines in gene expression and cis-regulatory element variation in Drosophila melanogaster. BMC Genomics 17, 981 (2016).

10. Karunakar P, Bhalla A, Sharma A. Transgenerational inheritance of cold temperature response in Drosophila. FEBS Lett. 593, 594–600 (2019).

11. Kolaczkowski B, Kern AD, Holloway AK, Begun DJ. Genomic differentiation between temperate and tropical Australian populations of Drosophila melanogaster. Genetics 187, 245–260 (2011).

12. Lesch, B. J. et al. Intergenerational epigenetic inheritance of cancer susceptibility in mammals. Elife doi: 10.7554/eLife.39380 (2019).

13. Li, J. et al. Genomic hypomethylation in the human germline associates with selective structural mutability in the human genome. PLoS Genet. 8, e1002692 (2012).

14. Lyne R, Smith R, Rutherford K, Wakeling M, Varley A, Guillier F, Janssens H, Ji W, Mclaren P, North P, Rana D, Riley T, Sullivan J, Watkins X, Woodbridge M, Lilley K, Russell S, Ashburner M, Mizuguchi K, Micklem G. FlyMine: an integrated database for Drosophila and Anopheles genomics. Genome Biol. 8, R129 (2007).

15. Mateo L, Rech GE, González J. Genome-wide patterns of local adaptation in Western European Drosophila melanogaster natural populations. Sci, Rep. 8, 16143 (2018).

16. Mohammad, F., Singh, P. & Sharma, A. A Drosophila systems model of pentylenetetrazole induced locomotor plasticity responsive to antiepileptic drugs. BMC Syst. Biol. 3, 11 (2009).

17. Molaro, A. et al. Sperm methylation profiles reveal features of epigenetic inheritance and evolution in primates. Cell 146, 1029–1041 (2011).

18. Perez, M. F. & Lehner, B. Intergenerational and transgenerational epigenetic inheritance in animals. Nat. Cell Biol. 21, 143–151 (2019).

19. Pool JE, Braun DT, Lack JB. Parallel Evolution of Cold Tolerance within Drosophila melanogaster. Mol. Biol. Evol. 34, 349–360 (2017).

20. Radford, E. J. Exploring the extent and scope of epigenetic inheritance. Nat. Rev. Endocrinol. 14, 345–355 (2018).

21. Ray M, Acharya S, Shambhavi S, Lakhotia SC. Over-expression of Hsp83 in grossly depleted hsrω lncRNA background causes synthetic lethality and l(2)gl phenocopy in Drosophila. J. Biosci. doi: 10.1007/s12038-019-9852-z (2019)

22. Rifkin SA, Houle D, Kim J, White KP. A mutation accumulation assay reveals a broad capacity for rapid evolution of gene expression. Nature 438, 220–223 (2005).

23. Sawarkar R, Sievers C, Paro R. Hsp90 globally targets paused RNA polymerase to regulate gene expression in response to environmental stimuli. Cell 149, 807–818 (2012).

24. Sawarkar R, Paro R.: an emerging hub of a networker. Trends Cell Biol. 23, 193–201 (2013).

25. Sharma, A. & Singh, P. Detection of transgenerational spermatogenic inheritance of adult male acquired CNS gene expression characteristics using a Drosophila systems model. PLoS One 4, e5763 (2009).

26. Sharma, A. Systems genomics analysis centered on epigenetic inheritance supports development of a unified theory of biology. J. Exp. Biol. 218, 3368–3373 (2015).

27. Sharma, A. Multi-omics data analysis implicating epigenetic inheritance in evolution and disease. BBA - General Subjects, https://doi.org/10.1016/j.bbagen.2019.129477 (2019).

28. Skvortsova, K., Iovino, N. & Bogdanović, O. Functions and mechanisms of epigenetic inheritance in animals. Nat. Rev. Mol. Cell Biol. 19, 774–790 (2018).

29. Teltumbade, M., Bhalla, A. & Sharma, A. Paternal inheritance of diet induced metabolic traits correlates with germline regulation of diet induced coding gene expression. Genomics doi: 10.1016/j.ygeno.2019.04.008 (2019).

30. von Heckel, K., Stephan, W. & Hutter, S. Canalization of gene expression is a major signature of regulatory cold adaptation in temperate Drosophila melanogaster. BMC Genomics 17, 574 (2016).

31. Yablonovitch AL, Fu J, Li K, Mahato S, Kang L, Rashkovetsky E, Korol AB, Tang H, Michalak P, Zelhof AC, Nevo E, Li JB. Regulation of gene expression and RNA editing in Drosophila adapting to divergent microclimates. Nat. Commun. 8, 1570 (2017).

32. Zare, A., Johansson, A. M., Karlsson, E., Delhomme, N. & Stenberg, P. The gut microbiome participates in transgenerational inheritance of low-temperature responses in Drosophila melanogaster. FEBS Lett 592, 4078–4086 (2018).

33. Zhao L, Wit J, Svetec N, Begun DJ. Parallel Gene Expression Differences between Low and High Latitude Populations of Drosophila melanogaster and D. simulans. PLoS Genet. 11, e1005184 (2015).

